# Alliance formation and complexity of Indo-Pacific bottlenose dolphins (*Tursiops aduncus*) around Mikura Island, Japan

**DOI:** 10.64898/2026.06.12.731994

**Authors:** Hibiki Nishitani, Tadamichi Morisaka, Kazunobu Kogi, Motoi Yoshioka

**Affiliations:** Graduate School of Bioresources, Mie University, Mie, Japan; Mikurashima Tourism Association

**Keywords:** alliance, cooperation, mammal, social behavior, toothed whale, *Tursiops*

## Abstract

In this study, we investigate alliance formation and complexity in male Indo-Pacific bottlenose dolphins around Mikura Island using five years of data collected through underwater observations. Focusing on 18 mature males, we examined affiliative behaviors (proximity and rubbing), consortships, and associations. To determine male relationships, we evaluated a simple model (small units) and a complex model (large units). In both models, units were identified by association, and models were evaluated by the extent to which affiliative behaviors and consortship were concentrated within units. All types of behaviors were more concentrated within units determined by the complex model than the simple model, and thus the former was further investigated. The unit sizes determined by the complex model were three, seven, and eight, and variation in association frequency was observed within units. Within units, two to four males engaged in a single consortship irrespective of association frequency. Given that units are mediated by affiliative and cooperative relationships, it is reasonable to interpret units as alliances. Considering the variation in both alliance size and within-alliance relationships, as well as the fact that only a few males cooperate in a single consortship, we suggest that a multi-level structure is plausible in the Mikura community.

## 1 Introduction

An alliance is a cooperative relationship in which individuals repeatedly join forces to compete against conspecifics (Harcourt and de Waal 1992). Currently, coalitional behavior or alliances have been documented in mammals across five orders (Primates, Artiodactyla, Perissodactyla, Proboscidea, and Carnivora) and over 60 species (e.g., Sterck et al. 1997, Olson and Blumstein 2009, Smith et al. 2010, Bissonnette et al. 2014, Bissonnette et al. 2015, Lukas and Clutton-Brock 2018, Smith et al. 2022, Smith et al. 2023). Given that alliances are regarded as one of the most complex social traits, cross-species comparisons of alliances in mammals are crucial for understanding the evolution of social behavior and cognitive abilities from both phylogenetic and adaptive perspectives (Harcourt and de Waal 1992).

In Cetacea, alliance formation to access for females has been reported in males of two species of the genus *Tursiops* (common bottlenose dolphin: *T. truncatus* and Indo-Pacific bottlenose dolphin: *T. aduncus*), with variations in formation (forming vs. non-forming) and complexity (uni-level or multi-level) within and across communities (Connor et al. 1992a,b, Möller et al. 2001, Wells 2014, Connor and Krützen 2015, Ermak et al. 2017, Connor et al. 2022, Brightwell and Gibson 2023). The society of *Tursiops* is characterized by fission-fusion dynamics, where group size and composition change temporally (Connor et al. 2000, Aureli et al. 2008). In this fluid society, males form alliances in the context of reproduction, such as mating with females, and defending or thieving females from other males or alliances (Connor et al. 2000). Genetic analyses have shown that alliance formation enhances reproductive success at the individual and/or alliance levels (Krützen et al. 2004, Wiszniewski et al. 2012b, Gerber et al. 2022, Duffield and Wells 2023). Thus, selection seems to favor alliance formation among males. Nevertheless, findings of alliances in *Tursiops* within and across regions showed remarkable variation in both alliance formation and complexity (Brightwell and Gibson 2023). Given the variation in alliances in *Tursiops*, it may contribute to identifying the factors that favor alliance formation and complexity through cross-community comparative analysis (Connor et al. 2000, Whitehead and Connor 2005, Möller 2012, Connor and King 2026).

To explain variation in male alliances, the effects of inter-male competition, body size, and sexual size dimorphism have been hypothesized (Connor et al. 2000, Whitehead and Connor 2005, Möller 2012, Connor and King 2026). The inter-male competition hypothesis has been tested by theoretical analyses, cross-community and within-community comparisons. Whitehead and Connor (2005) conducted agent-based simulations and showed that inter-male competition could explain variation in alliance size. Empirical studies have approximated inter-male competition as the communication range among individuals (hereafter, active space), community density, and the operational sex ratio. Brightwell and Gibson (2023) showed a positive relationship between community density and both alliance formation and complexity through cross-community comparisons. In Shark Bay, studies showed variation in alliances along environmental gradients within the same community (Connor et al. 2017, O’Brien et al. 2020, Sørensen et al. 2024). In contrast, hypotheses regarding male body size and sexual size dimorphism remain less well tested, both empirically and theoretically. Connor et al. (2000) and Connor and King (2026) suggested that when variation in male body size is large, males may be able to win competition without forming alliances, and when sexual size dimorphism is pronounced, males may gain access to females without cooperation. These factors are therefore expected to influence alliances.

Around Mikura Island, approximately 110 to 160 Indo-Pacific bottlenose dolphins (*Tursiops aduncus*) are resident year-round (Kogi et al. 2004, Sakai et al. 2025), and this community relatively exhibits distinct environmental and social characteristics compared to previously studied communities (Brightwell and Gibson 2023). Most communities examined in previous studies primarily inhabited bays or estuarine systems, whereas the Mikura community inhabits waters surrounding a pelagic island. Such differences may influence alliances through their effects on the distribution and abundance of prey and predators, which primarily influence female ecology and the active space among individuals (Connor et al. 2000, Connor and King 2026). In addition, the Mikura community shows low emigration and immigration, and a high population density, which affect male competition (Tsuji et al. 2017). Furthermore, compared with the well-studied *T*. *aduncus* community in Shark Bay, individuals in the Mikura population differ in body size (male asymptotic length is 201 cm in Shark Bay, van Aswegen et al. 2019; 256 cm in Mikura Island, Morisaka et al. 2022). Underwater photo- and video-identification surveys have been conducted in this population since 1994, resulting in the accumulation of extensive information on individual identities, community characteristics, and ecology (Sakai et al. 2025). Knowledge of alliance formation in the Mikura community, which possesses these distinctive ecological features and an extensive database, will play an important role in understanding the relation between environmental conditions and alliances through comparative analyses.

While previous studies conducted in the Mikura community have suggested the social relationships among males (Sakai et al. 2006, Sakai et al. 2010, Miyanishi et al. 2023), no data have been presented showing that these relationships are functionally equivalent to alliances reported in previous studies. In previous studies of the genus *Tursiops*, alliances are generally determined by a higher frequency of association, which is the proportion of time that a dyad spends together (Brightwell and Gibson 2023). Given the challenges of direct behavioral observations in free-ranging cetacean research, it is often unavoidable to determine social relationships or alliances using proxy measures such as association (Samuels and Tyack 2000, Whitehead 2008). However, Connor and King (2026) proposed that a relationship measured by association can be interpreted as an alliance when individuals associate regularly, maintain affiliative interactions, and there are specific functional explanations for the relationship (see Endnote 1). Given that research on the Mikura community is valuable within a comparative framework, it is therefore necessary to determine alliances using both behavioral and association analyses.

Multi-level alliances have currently been reported from only two regions (Shark Bay, Connor and Krützen 2015; St. Johns River, Ermak et al. 2017, Brightwell et al. 2025). However, most previous studies have focused on alliance formation rather than complexity (Brightwell and Gibson 2023). As a result, it remains unclear whether the complex, multi-level alliances reflect environmental specificity or simply a lack of research. Alliance complexity has been studied primarily in Shark Bay, and a systematic framework for multi-level alliances has been proposed in that community. In summary, alliance complexity in Shark Bay is characterized by three elements: discrete levels, functional differentiation between levels, and continuous variation within levels (Connor and King 2026). Alliances in Shark Bay are conceptualized as comprising three levels. Second-order alliances, typically formed by 4 to 14 males, maintain long-term associations over several decades and are considered the core social units. Within these second-order alliances, 2 to 3 males form first-order alliances, whose stability varies along a continuum from labile to stable. In addition, several first- or second-order alliances form third-order alliances. First-order alliances primarily function in maintaining consortships, while second- and third-order alliances play roles in inter-male conflict. Although multi-level alliances have been documented in Shark Bay, in other regions, such as *T. truncatus* in Sarasota Bay, alliances typically consist of two or three males (Wells 2014). Therefore, when inferring the alliance complexity or structure within a community, it is necessary to evaluate multiple models such as uni-and multi-level models, simultaneously.

To further our understanding of the relationship between environmental conditions and alliance structure, we investigate alliance formation and its complexity in the Mikura community. First, we analyze affiliative behaviors (proximity and flipper rubbing) and consortships as well as association among males. Second, behavioral and association data were integrated to distinguish the male relationships. Specifically, we tested whether the relationship among males in this community are better described by a small, simple unit model consisting of two to three males or by a large, complex unit model. In this analysis, we considered that units inferred by association that correspond to males that actually interact with one another would be most appropriate. After determining unit membership, we analyze the relation between behavior and association within units. In addition, a sensitivity test was conducted to evaluate the uncertainty in unit membership.

## 2 Methods

### 2-1. Research location and community

The study location was Mikura Island, which is a dormant volcanic island with a circumference of 16.4 km and an area of 20.6 km², Tokyo, Japan (33°52.5′N, 139°36.1′E). Water depth around the island ranges from 2 to 45 m within 300 m of the shoreline. The island is located in the Kuroshio Current, and the seafloor substrate is predominantly composed of rounded stones approximately 30 cm in diameter.

The species inhabiting Mikura Island is an Indo-Pacific bottlenose dolphin (*Tursiops aduncus*, Kakuda et al. 2002). The size of this community fluctuates between approximately 110 and 160 individuals, with a sex ratio of approximately 1:1 (Kogi et al. 2004). Because the annual re-identification rate of individuals exceeds 86 % (Kogi et al. 2004) and immigration and emigration rates are low (Tsuji et al. 2017), this community is considered to have limited connection to neighboring communities.

### 2-2. Data collection and curation

Field surveys have been conducted on Mikura Island since 1994, from May to November (Sakai et al. 2025). Surveys were mainly carried out aboard tourist vessels, while a research vessel was occasionally used. The study area was primarily within 300 m of the coastline, and depending on sea conditions, either the entire island or only part of it was surveyed. Surveys were conducted one to three times per day, during the morning (07:00-12:00) and afternoon (13:00-17:00). Each survey usually lasted approximately two hours. A single field survey was conducted by one to four researchers. When a dolphin group was found at sea, researchers recorded information such as time, GPS location, and sea conditions. Then, researchers entered the water and recorded underwater video while snorkeling. Recording was conducted using video camera (SONY HDR-CX430, SONY HDR-XR550V, SONY HDR-SR12) placed in an underwater housing. When dolphins were no longer observable underwater, researchers returned to the boat and recorded the group information such as size and behavioral state. This sequence of actions, finding the dolphins, entering the water, recording video, and returning to the boat, was defined as an “encounter.” During a single field survey, between one and eight encounters occurred.

Collected data were used for individual identification following Kogi et al. (2004). Underwater video data and information on individuals and groups have been archived since 1994, and these data have been used for subsequent analyses.

### 2-3. Object individuals

In this study, we used data collected through individual identification surveys between 2015 and 2019. The males included in the analysis were 18 individuals that met the following criteria: 1) they were at least 15 years old in 2015, and 2) they were identified in every year from 2015 to 2019. The age threshold of 15 years was set to include only individuals that had reached sexual maturity (Kemper et al. 2014). The requirement for continuous identification was applied to avoid potential bias in association analyses caused by immigration or emigration (Whitehead 2008).

### 2-4. Behavioral analysis

We described behaviors previously reported among allied males based on previous studies (e.g., Connor et al. 1992a,b, Connor et al. 2006, Connor and Krützen 2015). As affiliative behaviors among males, we recorded proximity and flipper rubbing. Proximity was defined as “swimming in the same direction while another individual is within 0.5 body lengths, two body widths, and two body heights,” using the dolphin’s body as a reference unit. When recording proximity, a chain rule was applied. When individual A was in close proximity to individual B and individual B was in close proximity to individual C, individuals A and C were also considered to be in proximity. Rubbing was defined following Sakai et al. (2006) as “one dolphin contacts another dolphin with its pectoral fin (flipper) and either or both dolphins actively move the touching body parts back and forth.”

We also recorded behaviors directed from males toward females, as well as agonistic interactions among males. The behaviors of allied males in this species have been described primarily in Shark Bay (Connor and Krützen 2015) and anecdotal and verbal descriptions of these behaviors have also been reported from other regions (e.g., Félix 1997, Möller et al. 2001, Parsons et al. 2003, Brightwell et al. 2025). In the present study, we recorded the following behaviors based on descriptions from Shark Bay studies: “capture,” “bolt,” “aggression,” and “inter-male aggression.” Capture consisted of behavioral elements such as formation swimming, in which males position themselves behind and on either side of the female, as well as rapid chases directed at the female. Bolt refers to the female behavior during capture, in which the female initiates rapid swimming to escape or attempt to escape from capture. Aggression includes indirect displays such as jaw claps as well as direct behaviors such as biting and tackling. Inter-male aggression refers to aggressive interactions occurring between males. Although pop sounds are included in the alliance criteria (Connor and Krützen 2015, Connor et al. 2022), their definition requires detailed behavioral and acoustic data. Therefore, pop sounds were not included in this study. These behaviors were recorded using daily 1–0 sampling.

We also recorded the reproductive status of females involved in consortships and whether they gave birth in the following year. The reproductive status of a female during consortship was classified as “receptive” when one of the following conditions was met: 1) when the female was not associated with a calf, she had either given birth at least once or was ≥9 years old, 2) when the female was associated with a calf, the calf was ≥2 years old, or 3) when the female gave birth in the year following the consortship. Condition 1 was determined based on the mean age at first reproduction in this community (unpublished data from Mikurashima Tourism Association). Condition 2 was based on the average interbirth interval in this community, which suggests that females could become reproductively receptive when their calf is approximately 2 years old (Kogi et al. 2004). Reproductive status was recorded as “unknown” in the following cases: 1) when a female involved in a consortship was not identified in the following year, or 2) when no birth had been recorded for more than 6 years since the female’s last observed birth. This criterion was based on the average interbirth interval in this community (Kogi et al. 2004) and the possibility of physiological reproductive cessation in older females. Similarly, whether the female gave birth was recorded as “unknown” when the female involved in the consortship was not identified during following year.

### 2-5. Association analysis

Association analyses were conducted to characterize association patterns at the community level and to determine alliance formation and complexity. In this study, associations were defined using the “gambit of the group” assumption, in which all individuals belonging to the same group are considered to be associated (Whitehead 2008). A group was defined following Morisaka et al. (2023). Under this definition, groups were determined using the following procedure: 1) each encounter was initially defined as a group, and 2) when the same individual was identified across multiple encounters within the same survey, those encounters were merged. Associations were recorded using daily 1–0 sampling. Associations were quantified using the simple ratio index and the half-weight index (hereafter SRI and HWI, respectively; Cairns and Schwager 1987, Croft et al. 2008, Whitehead 2008, Farine and Whitehead 2015, Hoppitt and Farine 2018). These indices range from 0 to 1, where 0 indicates that two individuals were never identified in the same group, and 1 indicates that two individuals were always identified in the same group. These indices were calculated using pooled data from the five-year study period.

Because the results obtained using SRI and HWI were consistent, only the results based on SRI are presented in Result section (see Supplemental Material).

### 2-6. Statistical analysis

All analyses were conducted in R version 4.5.2 (R Core Team 2025). The analyses were performed with reference to SOCPROG 2.10 uncompiled version (Whitehead 2009).

To examine whether individuals showed non-random association, we conducted a permutation test. The null hypothesis was that the standard deviation (SD) of the observed association indices was equal to the expected SD, which was calculated from Monte Carlo simulations. To generate associations expected under random association, permutations were performed by swapping association records only within the same sampling day (Whitehead 2008). A swap involves selecting two dyads from the association matrix and exchanging their association records. The number of permutations was set to 20,000, and the number of swaps per permutation was set to 1,000.

We also calculated social differentiation (s) as a parameter describing the nature of the association indices (Whitehead 2008). This parameter is defined as the estimated coefficient of variation of the true association indices and reflects the variability of association indices within a community. To estimate social differentiation, we first estimated the true association indices, which represents the proportion of the time that a dyad was actually associated. When the parameter is close to 0, relationships within the community are homogeneous, whereas s near or greater than 1 indicates heterogeneous relationships. In this study, parameter s was estimated using maximum likelihood methods, and standard errors (SE) were estimated using a bootstrap method with 1,000 resamples.

We further evaluated the precision and accuracy of the calculated association indices (Whitehead 2008). SE of the association indices was estimated using a bootstrap method with 1,000 resamples. The accuracy of the association indices was estimated by the correlation coefficient (r) between the true association indices and the observed association indices. The r ranges from 0 to 1, and the value greater than 0.8 are considered desirable. As with social differentiation, the true association indices were estimated using maximum likelihood methods, and their SE was estimated using a bootstrap method with 1,000 resamples.

To determine the alliance structure, we applied two association-based models that have been used in previous studies (e.g., Connor et al. 1992a,b, Möller et al. 2001, Owen et al. 2002, Ermak et al. 2017, Baker et al. 2020). One was called the “simple model,” in which units are defined as dyads that show the highest mutual association index with each other. However, trios are formed for individuals that do not have a mutual highest associate when the individual’s first and second most frequent associates both have that individual as their own second most frequent associate. Quartets or larger units may also be formed when ties occur. The second model was the “complex model.” In this model, units were determined using the following procedure. First, a dendrogram was constructed using association with the average-linkage method, and the dendrogram was evaluated using the cophenetic correlation coefficient (CCC). The CCC is a measure that evaluates how faithfully the generated dendrogram represents the relationships in the original dataset. CCC ranges from 0 to 1, and values greater than 0.8 are generally considered desirable. From the resulting dendrogram, modularity (Q) was calculated. Modularity is defined as the difference between the proportion of total association occurring within units (or clusters) and the expected proportion (Newman 2004, Whitehead 2008). Q was calculated using the following equation.

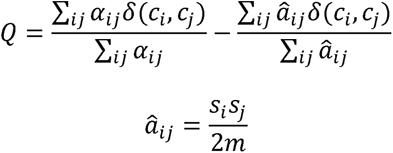

Here, α_*ij*_ represents the observed association index for dyad *ij*, and 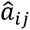 represents the expected association index if dyad *ij* associates randomly. *δ*(c_*i*_, c_*j*_ ) takes the value 1 when individuals *i* and *j* belong to the same cluster and 0 when they belong to different units. s_*i*_ is the sum of association indices for individual *i* (strength or gregariousness), and *m* is defined as ∑(*s*/2). *Q* ranges from 0 to 1, where 0 indicates that units are randomly arranged and 1 indicates that no associations occur between units. According to Newman (2004), units are considered to be clear when *Q* exceeds 0.3. In this study, modularity was calculated for each bifurcation of the dendrogram, and clusters were determined at the point where *Q* reached its maximum (*Q*_*max*_) (Lusseau 2007, Whitehead 2008).

To evaluate which model best represents actual relationships among males, we quantified the frequency of observed behaviors within and between units. We calculated the “within-unit interaction ratio (hereafter, IR),” defined as the frequency of interactions observed within alliances divided by the total frequency of interactions (within-unit + between-unit). Here, we assumed that alliances are mutually exclusive relationships and therefore expected that interactions would be concentrated within units and rarely observed between units. Based on this assumption, the model with the higher within-unit interaction ratio was considered to better represent the alliance structure.

After determining the alliance structure, we analyzed the relation between interactions and association within alliances (units) to infer internal alliance structure. Because the within-unit interaction ratio supported the cluster model (see Results), this analysis was conducted only for the results obtained from the complex model.

To quantify the uncertainty in the assignment of alliance membership, we conducted a sensitivity analysis using a jackknife simulation. In this analysis, two days of data were removed from each year (a total of 10 days), and a network was reconstructed from the remaining data. Cluster analysis was then performed to obtain alliance membership. This procedure was repeated 1,000 times. For each trial, we recorded whether each dyad *ij* exhibited the same unit membership as in the full dataset (0 = different, 1 = same). The consistency indices were calculated, and these ranged from 0 to 1, with values closer to 1 indicating that unit membership of each dyad *ij* had low uncertainty.

## 3 Results

### 3-1. Research effort

We conducted 283 days of surveys and analyzed 221 days of data on which at least one focal individual was identified between 2015 and 2019. The mean number of identification days per individual was 49.1 (SD = 18.9, range = 22 – 92). Detailed information is listed in the Supplemental Materials.

### 3-2. Behavior analysis

Proximity was observed 174 times across 53 dyads, flipper rubbing was observed 17 times across 15 dyads, and consortship was observed 19 times across 18 dyads. Table S3 shows details of consortship. Single consortships involved two males in 16 cases, three males in one case, and four males in one case. Most consortship events involved capture, especially formation swimming (n = 17), whereas capture, bolt, and aggression were observed in one case (Figure 1; see Supplemental Video). No aggressive interactions among males were observed. Twelve of 15 females (80%) were in a receptive state at the time of the consortship. Of these reproductively receptive females, 6 of the 12 (50%) gave birth in the following year.

**Figure 1.**
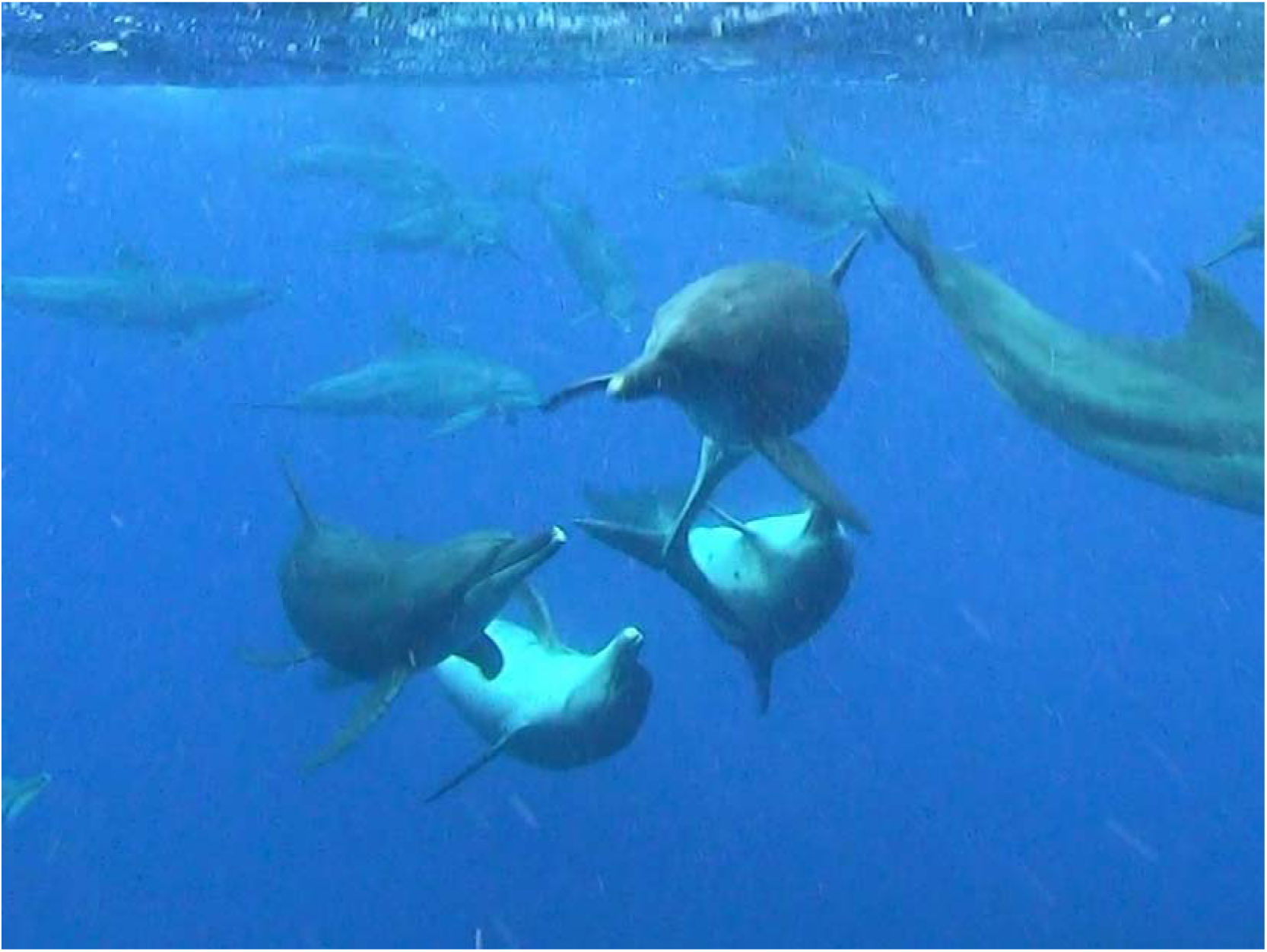
Aggressive consortship observed in the Mikura community. Four males were involved in this event (one male is outside the frame). In this event, immature and mature males surrounded the consorting group of four males and one female. See Supplemental Video.

### 3-3. Association analysis

The distribution of SRI was right-skewed (Figure S1). The permutation test showed that the SD of the observed SRI values was significantly greater than the expected SD (Table S4, Figure S1, one-sided test, p < 0.05). The social differentiation and correlation coefficient were estimated at 0.76 (SE = 0.06) and 0.86 (SE = 0.03), respectively (Table S4). The mean SE of the SRI was 0.03 (SD = 0.01, Figure S1).

### 3-4. Model analysis

Although all types of behavior were frequently observed between units determined by the simple model, these behaviors were concentrated within units determined by the complex model. The simple model divided individuals into five clusters, with cluster sizes of 2 in three units and of 3 in two units (Figure 2). The complex model divided individuals into three units, with sizes of 7, 3, and 8 (Figure 2, Figure 3). The CCC of the dendrogram was 0.96, and *Q_max_* was 0.33 (Figure 3). Under the simple model (Figure 4a,b,c), IR was 0.17 (9/53) for proximity, 0.40 (6/15) for rubbing, and 0.33 (6/18) for consortship. Under the complex model (Figure 4d,e,f), IR was 0.75 (40/53) for proximity, 0.87 (13/15) for rubbing, and 0.83 (15/18) for consortship.

**Figure 2.**
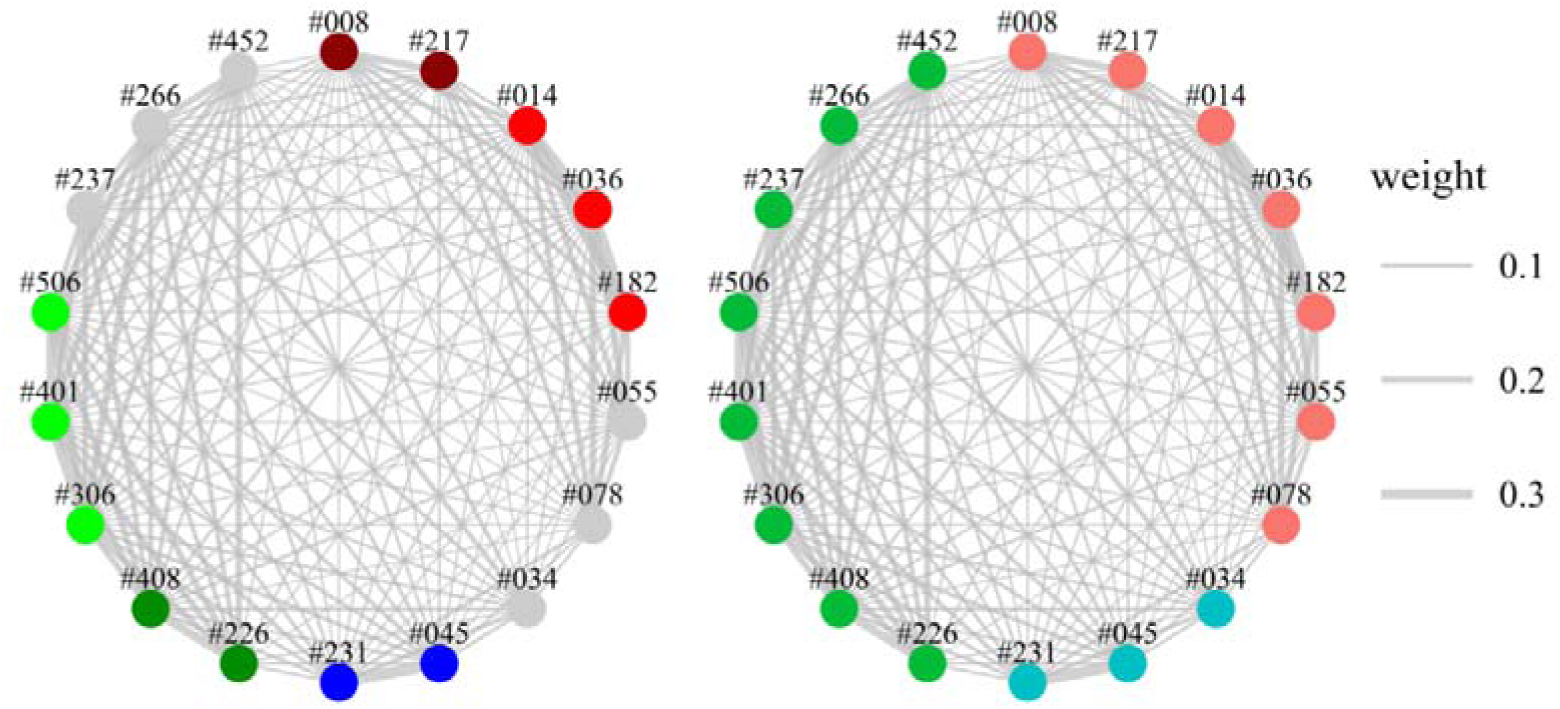
Sociogram with simple and complex model results (left = simple model, right = complex model). Nodes represent individuals and their color represents unit membership. Note that gray nodes represent individuals that were not assigned any units. Edges represent association and their width represents association indices.

**Figure 3.**
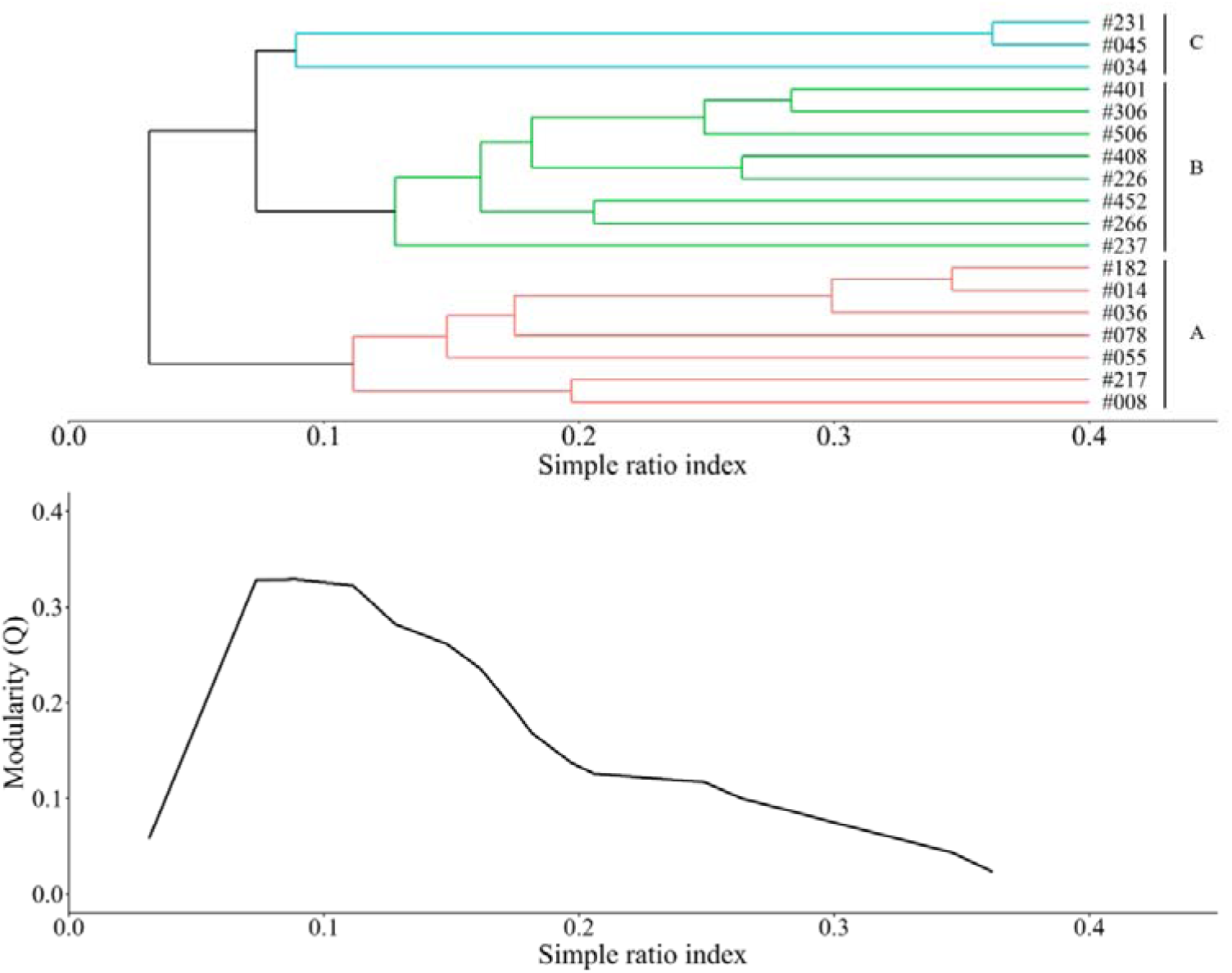
Dendrogram constructed by the average-linkage method. Colors represent units determined by the modularity method. “A”, “B” and “C” are convenient unit names. The graph below shows changes in modularity with dendrogram bifurcations.

**Figure 4.**
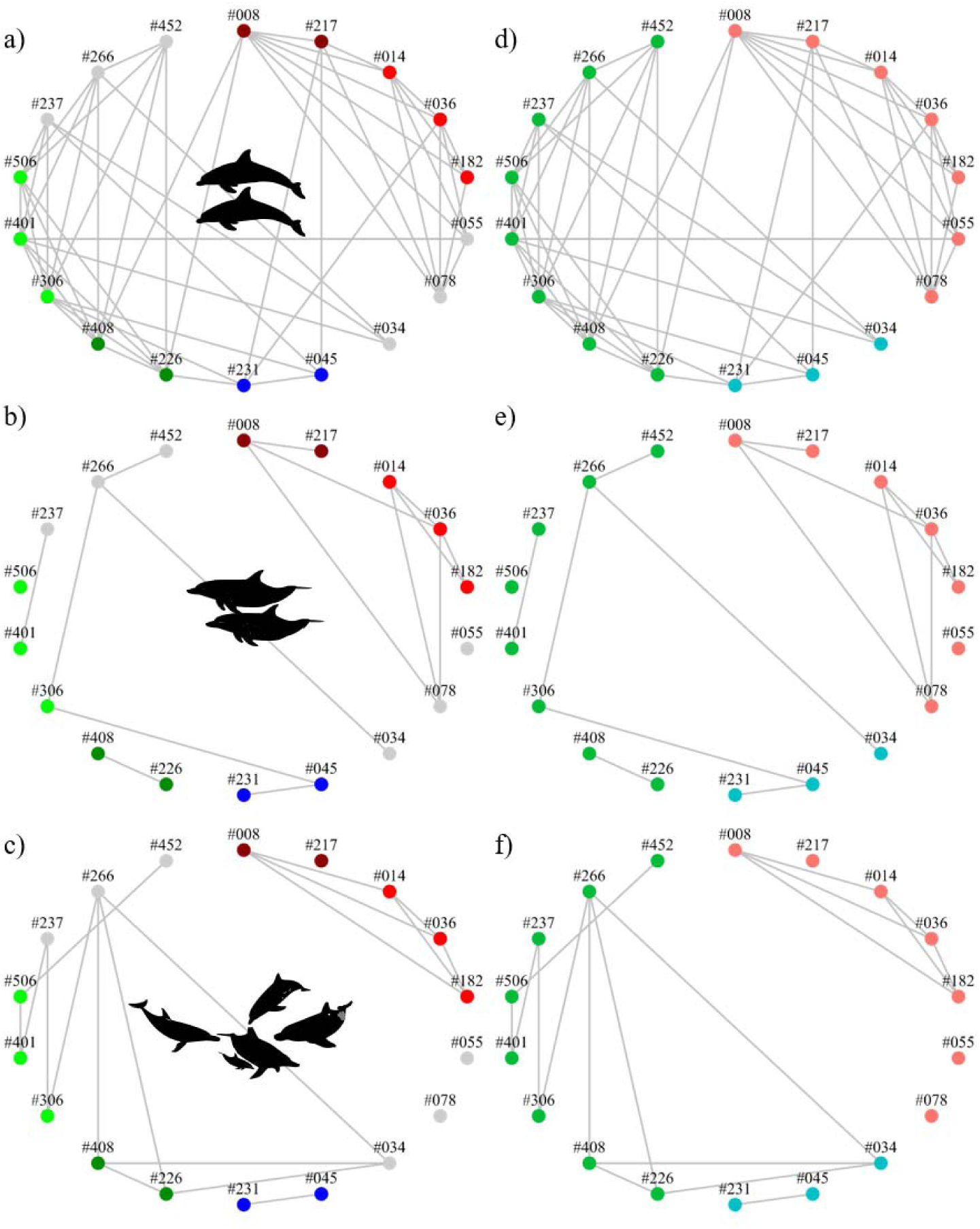
Behavioral observation within units determined by the simple (left) and complex models (right). Edges represent proximity (a,d), flipper rubbing (b,e), and consortship (c,f). Other attributes are the same as those in Figure 2.

### 3-5. Relation between behavior and association within cluster

All types of behavior were observed among males within units determined by the complex model, with a wide range of association frequencies (Figure 5). Within unit A, the mean SRI was 0.17 for proximity (SD = 0.08, range = 0.08 – 0.35), 0.22 for rubbing (SD = 0.09, range = 0.10 – 0.35), and 0.22 for consortship (SD = 0.10, range = 0.13 – 0.35). Within unit B, the mean SRI was 0.19 for proximity (SD = 0.04, range = 0.12 – 0.28), 0.21 for rubbing (SD = 0.04, range = 0.15 – 0.26), and 0.19 for consortship (SD = 0.05, range = 0.13 - 0.26). Within unit C, the SRI of the dyad in which proximity, rubbing, and consortship were observed was 0.36 (only one dyad was included).

**Figure 5.**
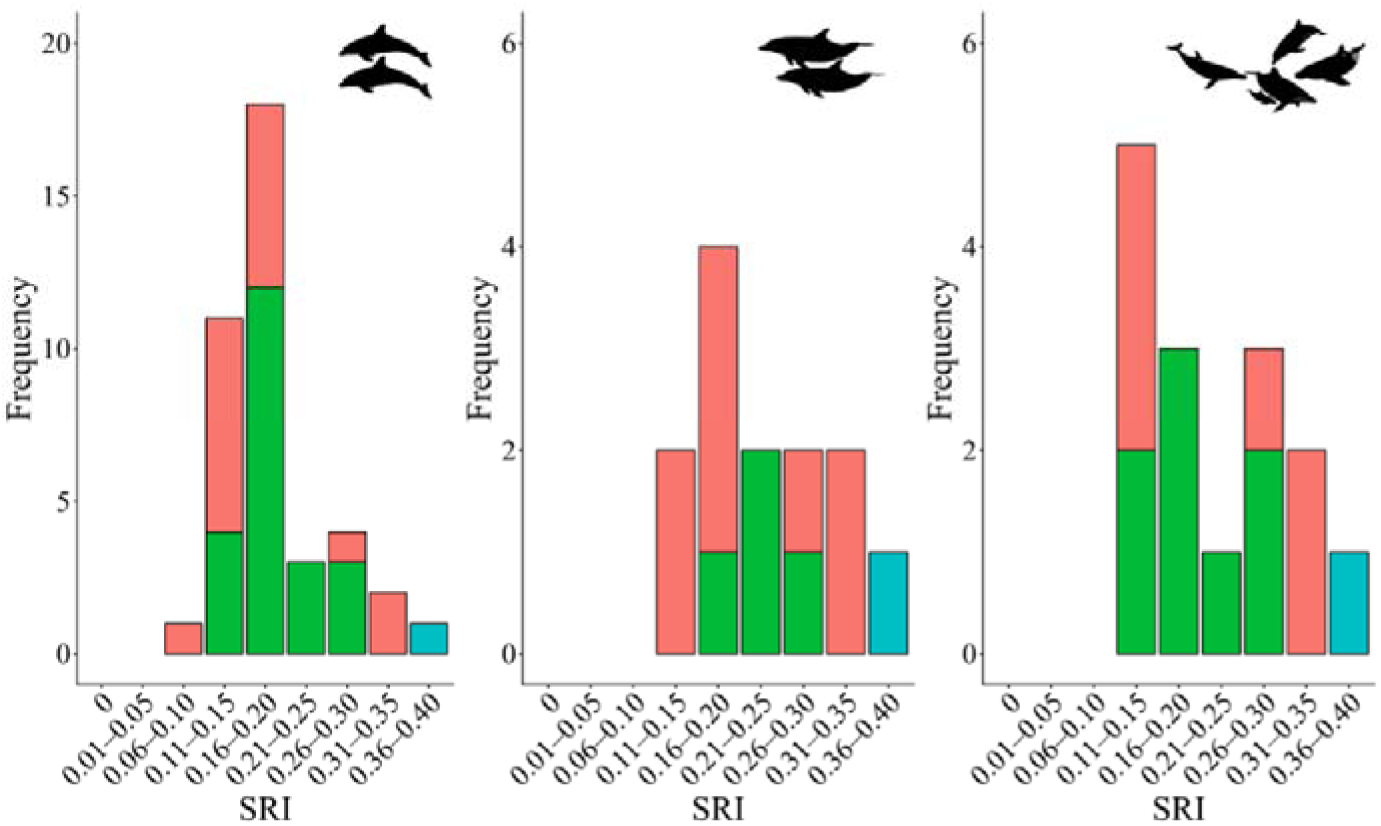
Within unit relation between SRI and behavior. The bar graphs, from left to right, show the results for proximity, rubbing, and consortship. Bar color represents units determined by the complex model (See Figure 2 and 3).

### 3-6. Sensitivity analysis

The jackknife simulation showed a high consistency index, but some uncertainty was observed. Simulations showed a mean consistency index of 0.97 (SD = 0.08, range = 0.66 – 1.00, Figure 6). Membership of unit A was completely stable (consistency index = 100%), whereas the membership of unit B and unit C varied across simulations (Figure 7). Specifically, in some simulations, unit B and C were merged into a single unit, whereas in others only individual #034 was assigned to unit C. However, individuals #045 and #231, and #226, #237, #266, #306, #401, #408, #452, and #506 always belonged to the same unit. The consistency index of #034 in unit C was 0.66 on average, whereas it was 0.79 in unit B on average. When the unit membership of #034 was treated as unknown, IR was 0.80 (40/50) for proximity, 0.93 (13/14) for rubbing, and 1.00 (15/15) for consortship. Within unit, the mean SRI of dyads was 0.19 (SD = 0.06, range = 0.08 – 0.36) for proximity, 0.23 (SD = 0.08, range = 0.10 – 0.36) for rubbing, and 0.21 (SD = 0.08, range = 0.13 – 0.36) for consortship.

**Figure 6.**
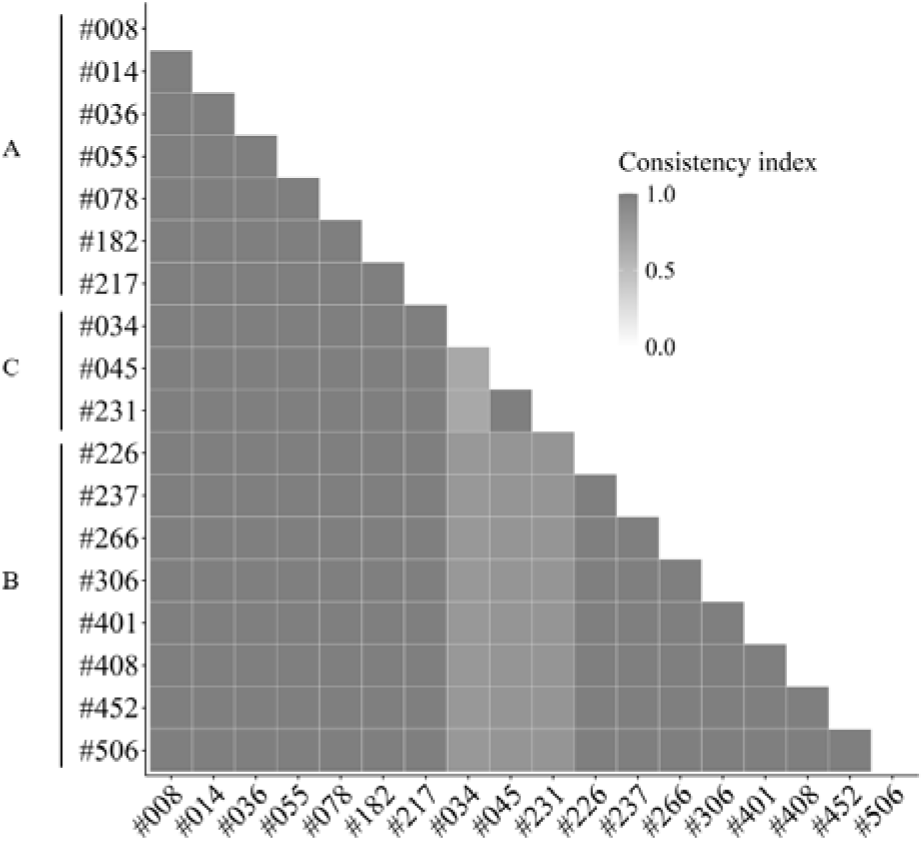
Consistency indices calculated by the jackknife simulation.

**Figure 7.**
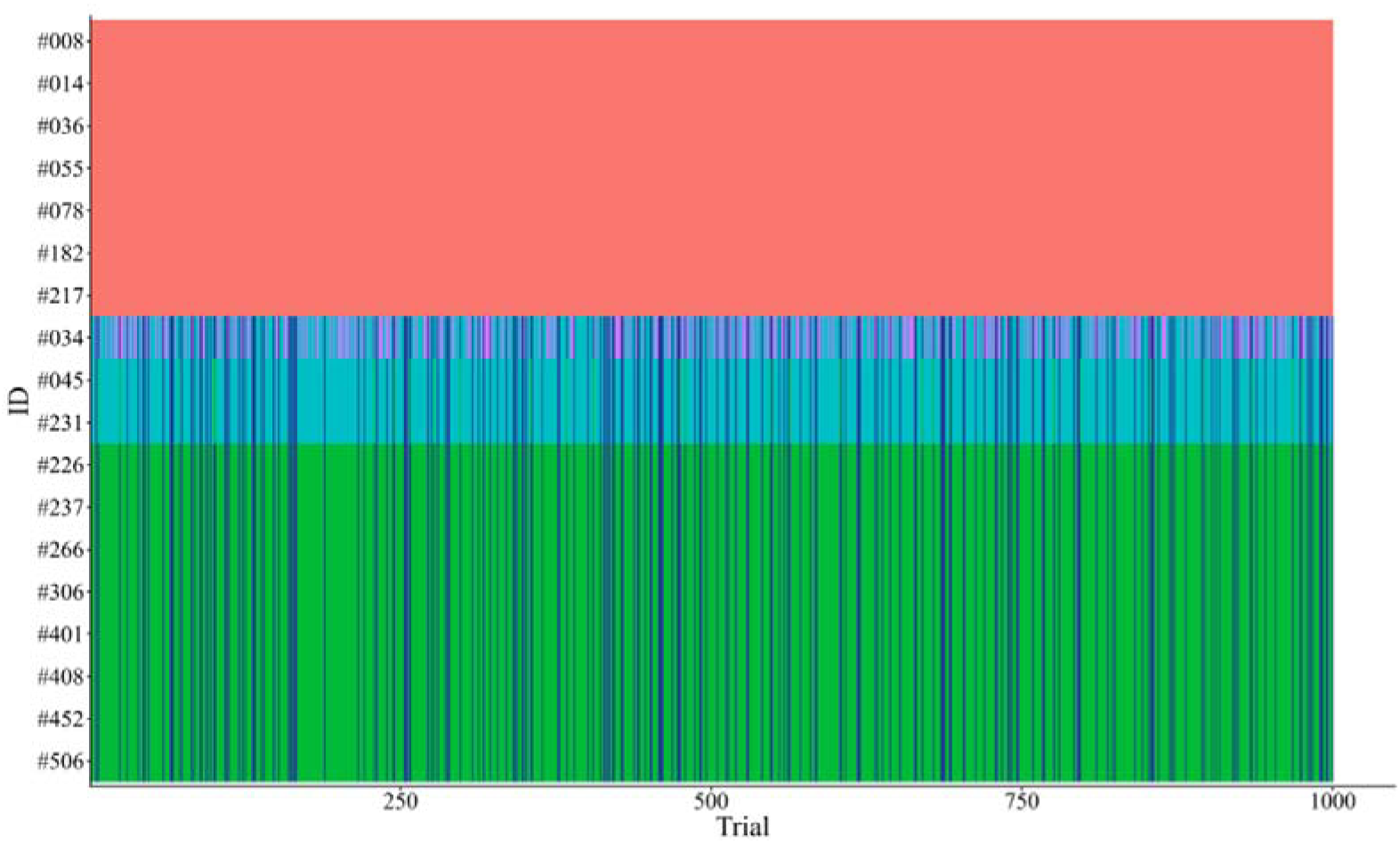
Details of the jackknife simulation. The x-axis represents 1,000 trials. Each color represents unit membership.

## 4 Discussion

This is the first study to simultaneously analyze both behavior and associations among mature male Indo-Pacific bottlenose dolphins around Mikura Island to determine alliance formation and complexity. Affiliative behaviors and consortship toward females were observed among males, and a single consortship event generally involved two males. In addition, the results of association analysis indicated that males in this community form non-random, heterogeneous social relationships. For all types of behaviors, such as proximity, flipper rubbing, and consortship, the within-unit interaction ratio (IR) was higher when units were determined using the complex model than simple model. This indicated that the units identified by the complex model more accurately reflected actual interaction patterns among males. Given that units determined by association frequency are mediated by affiliative and cooperative relationships, they satisfy the criteria of Connor and King (2026), and it is therefore reasonable to interpret these units as alliances. The alliance sizes determined by the complex model were three, seven, and eight males, and variation in association indices was confirmed within units. Considering that alliance sizes were relatively large and that few males engaged with a single consortships, we suggest that alliances in this community are best described by a multi-level structure like that observed in Shark Bay (Connor and King 2026).

### 4-1. Methodological differences and challenges

Our study was conducted using underwater observations, and the sampling method differed substantially from previous studies, which may cause some quantitative differences. The association indices are influenced by identification rate per group (Whitehead 2008) and group definition (Syme et al. 2022). While relatively high thresholds of association indices (e.g., 0.5 or 0.8) have been used to determine alliances in previous studies (e.g., Owen et al. 2002, Ermak et al. 2017), the maximum SRI recorded in this study was 0.36 (0.51 for HWI, see Supplemental Materials). The relatively small association indices in the Mikura community are likely due to the lower identification rate per group during underwater sampling (Morisaka et al. 2023). Importantly, however, even when the association indices are relatively small, association can still be used to distinguish individuals that actually interact with one another. The units determined by association showed high within-cluster interaction ratios and analyses conducted using different association indices such as HWI produced the same results, indicating the robustness of our findings (see Supplemental Materials).

In addition, the group definition used in this study is compatible with methods used in boat-based observations (Morisaka et al. 2023).

Although the within-cluster interaction ratio used in this study could evaluate false negatives, it does not allow the evaluation of false positives. For example, it can show that males that actually engage in consortships with one another are rarely assigned to different units. However, it cannot show whether some males within a unit are not truly allied. This limitation arises because behavioral observations are relatively rare, making it difficult to distinguish whether a dyad that belonged to the same unit truly does not interact or whether the interaction was simply missed. As a result, evaluating false positives is inherently difficult. Nevertheless, considering that the association indices within units were not strongly related to consortship observations, and that consortship was not observed between units, it is therefore reasonable to interpret the units identified by the complex model as alliances. In the future, more detailed behavioral observations (e.g., Connor et al. 1992a,b, Connor and Krützen 2015, Hill-Cousins 2025), analyses of developmental processes (e.g., Gerber et al. 2021, Gerber et al. 2022), genetic assessment of reproductive success (Krützen et al. 2004, Wiszniewski et al. 2012b, Gerber et al. 2022, Duffield and Wells 2023), and acoustic or cognitive analyses (e.g., Connor and Smolker 1996, King et al. 2018, King et al. 2021, Chereskin et al. 2022, Chereskin et al. 2024) will likely provide further insights into the structure of alliances.

### 4-2. Alliance complexity in Mikura community

Based on the result of complex model, we propose the structure of alliances in the Mikura community illustrated in Figure 8. The units identified by the complex model appear to be mediated by affiliative interactions and are capable of cooperating within units regardless of association strength. During a single consortship, two to four males cooperated. This structure satisfies two of the three conditions proposed for multi-level alliances in Shark Bay (discrete levels and variation within levels, Connor and King 2026). Although this suggests a multi-level structure, it is unclear whether different levels have distinct functions. In Shark Bay, the functions of higher-order alliances (e.g., second- and third-order alliances) are considered to involve inter-male and inter-alliance competition (Connor et al. 1992a,b, Connor et al. 2011, Connor and Krützen 2015, Connor et al. 2022). However, no agonistic interactions among males or alliances were observed in this study. Consequently, the functional differences among levels remain unclear, and further research is required.

**Figure 8.**
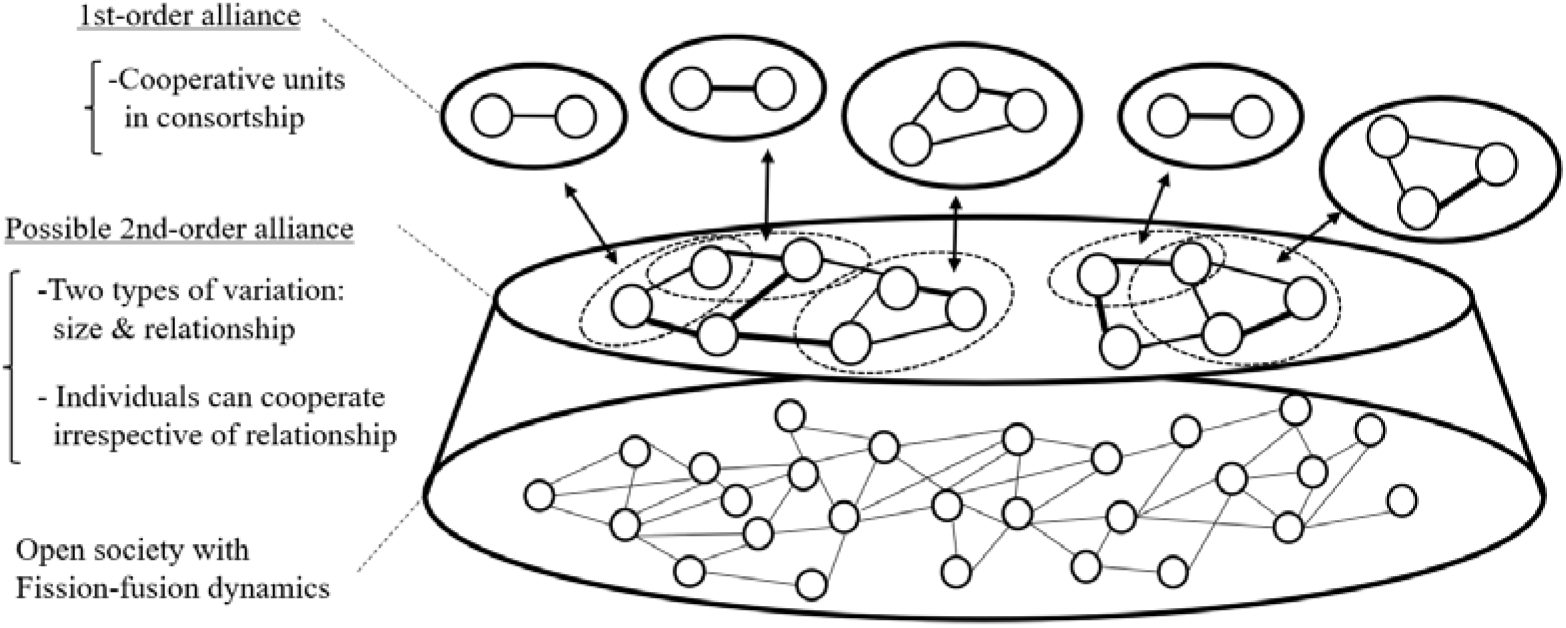
Schematic model of alliance structure in the Mikura community.

There is uncertainty in unit membership, particularly regarding the membership of #034, the relationship between unit B and unit C, and unit size. Two hypotheses may explain the uncertainty in the membership of #034. One is that #034 belongs to unit B. Indeed, affiliative behaviors and consortship involving #034 were observed with members of unit B, and the consistency index for assignment to unit B was higher than unit C. The alternative hypothesis is that #034 switched alliance membership during 5-years study period or belonged to multiple alliances. Theoretical work has considered alliance switching (Whitehead and Connor 2005) and analyses that account for overlapping memberships have not yet been conducted. Indeed, the current complex model (cluster analysis) assumes that each individual belongs to only one unit. Future analyses incorporating alliance switching or multiple memberships may provide important insights into alliance structure.

Regarding unit B and unit C, some simulations treated them as a single cluster. However, except for #034, individuals within both units were always assigned to the same unit. Although consortship was not observed between units, affiliative behaviors were occasionally observed. This pattern may be similar to the third-order alliance in Shark Bay (Freedman et al. 2023) and suggests that the unit B and unit C may form a third-order alliance. This hypothesis requires rigorous testing through further behavioral observations and cognitive experiments (King et al. 2021, Connor et al. 2022, Connor and King 2026). Finally, because focal males were filtered by age and identification history, alliance size may have been underestimated. In particular, individuals aged 10-14 years, representing a transitional reproductive stage, were excluded from the analysis. If these individuals belong to the units identified in this study, the true alliance size may be underestimated.

### 4-3. Geographical variation in alliances

The inter-male competition hypothesis is regarded as a leading explanation for variation in alliances, and male competition is typically approximated by factors such as active space, community density, and the operational sex ratio (Connor et al. 2000). Habitat characteristics are expected to influence active space, with more open environments assumed to allow larger active spaces, thereby increasing encounter rates among males (Connor et al. 2000, Brightwell and Gibson 2023). Although the Mikura community inhabits waters around a pelagic island and may appear to occupy an open environment, our surveys were restricted to within 300 *m* of the coastline, and the ranging patterns of this community remain unknown. Therefore, it is difficult to assume an active space based on habitat characteristics and to position the results of this study within this framework. Brightwell and Gibson (2023) suggested that community density provides some support for predictions of alliance formation and complexity, whereas the operational sex ratio, estimated using mean interbirth interval, shows weaker support. Although the estimated density of the Mikura community may be overestimated due to the limited survey area, it is relatively high (6.55 individuals/km²) compared with other studied communities, which is consistent with theoretical predictions and cross-community analysis (Connor and Whitehead 2005, Brightwell and Gibson 2023). With respect to the operational sex ratio, the number of mothers with a calf divided by the number of all mature females ranged from 0.64 to 0.91 in the Mikura community, indicating a male-biased operational sex ratio (Kogi et al. 2004). However, because few studies have estimated the operational sex ratio in this way, it is difficult to place these values in a comparative context. In addition, the mean interbirth interval in this community was 3.4 years (Kogi et al. 2004), which is similar to those reported in other communities (Brightwell and Gibson 2023). Therefore, the operational sex ratio in this community does not provide clear support for the hypothesis.

Body size, including total length and sexual size dimorphism, has been proposed to influence variation in alliances. In Shark Bay and Mikura Island, both of which assume multi-level alliances, the difference in asymptotic length is approximately 50 cm (201 cm at Shark Bay, van Aswegen et al. 2019; 256 cm at Mikura Island, Morisaka et al. 2022). Connor and King (2026) assumed that as mean body size increases, variance in body size also increases, and that this variance affects differences in male competitive ability. Variation in competitive ability among males may, in turn, influence the benefits of alliance formation. The multi-level alliances in two communities with different body sizes may suggest that body size does not strongly influence alliance complexity. Alternatively, it may indicate that a difference of approximately 50 cm in mean body size is not sufficient to influence alliance formation. Sexual size dimorphism (male/female) is 1.01 in Shark Bay (van Aswegen et al. 2019) and 1.04 in Mikura Island (Morisaka et al. 2022), indicating little difference between the two communities. Because few studies have estimated sexual size dimorphism, it is difficult to place these values in a comparative context. However, it appears that both communities show small sexual size dimorphism. It should be noted that the above comparisons are based on body length rather than body mass. In addition, it remains unclear whether body length and mass themselves accurately reflect male competitive ability in an aquatic, weak gravity environment. Instead, traits such as maneuverability may play a more important role in inter-male competition and access to females (Connor et al. 2000).

At a broad habitat scale, alliances have been reported not only in bays and estuarine systems but also in waters around pelagic islands (Brightwell and Gibson 2023). In addition, multi-level alliances have been documented in both bay and estuarine systems (Shark Bay, Connor and Krützen 2015; St. Johns River, Ermak et al. 2017, Brightwell et al. 2025) and around pelagic islands (this study). This suggests that macro-scale habitat itself may not be a useful predictor of variation in alliance structure. In contrast, within-community variation in alliance structure associated with environmental gradients has been reported in Shark Bay (Connor et al. 2017, O’Brien et al. 2020, Sørensen et al. 2024). Therefore, in addition to macro-scale comparisons, incorporating a finer-scale perspective is likely important for understanding the drivers of variation in alliances. It should be noted that comparative evaluation of alliance complexity is often difficult. Except for a limited number of studies (e.g., Ermak et al. 2017, Brightwell et al. 2020), most previous studies have not explicitly examined multi-level alliance structures. Therefore, the absence of reported multi-level alliances in those regions does not necessarily indicate that such structures are absent.

## 5 Conclusion

This is the first study to investigate alliance formation and complexity in the Mikura Island community. By combining behavioral observations and association analyses, we showed that males in this community form alliances. We also showed that a multi-level structure is more plausible for this community. Our long-term individual identification research has generated individual, behavioral, and genetic data (Sakai et al. 2025). Furthermore, our study employs detailed underwater observations, a method that is not feasible at many other study sites. By integrating these data and methods with the unique environmental and ecological characteristics of this community and conducting comparative analyses, future research will contribute to clarifying both the diversity and the ubiquity of alliances.

## Endnote

1. Nishitani H and Morisaka T (2026). *bioRxiv*: doi.org/10.64898/2026.01.14.699404; This article is currently under review. It points out that the association criteria used to define alliance formation in previous studies are inconsistent, which has prevented researchers from determining whether variation in alliances reflects inherent population-level characteristics or pseudo-variation arising from differences in criteria. Through a systematic literature review, this study suggests that this issue remains unresolved and provides empirical support for the suggestion of Connor and King (2026).

## Acknowledgment

This study was made possible by the financial and logistical support of the Mikurashima Tourism Association. We thank the dolphin-watching tour captains and guides, members of the identification research team (mido) and local villagers for supporting our research. We are grateful to the late Dr. Kyoichi Mori at Teikyo University of Science for his insightful perspective on cetacean biology, and to Dr. Masaki Shimada at Teikyo University of Science for his valuable advice on analytical methods. We appreciate Takuya Aoki for drawing the illustrations to use in the figures.

