## Supplemental data for "Alliance formation and complexity of Indo-Pacific bottlenose dolphins (*Tursiops aduncus*) around Mikura Island, Japan"

**Supplemental Materials**

Figure 4 in the main text is identical regardless of whether simple ratio index (SRI) or half weight index (HWI) is used. Therefore, the version of Figure 4 generated using HWI is not shown here.

Table S1. Survey days on which at least one object male was identified in each of the 5 years.

| Year | Survey days |
| --- | --- |
| 2015 | 43 |
| 2016 | 50 |
| 2017 | 44 |
| 2018 | 43 |
| 2019 | 41 |
| Total | 221 |

Table S2. Number of identifications of each individual in 5 years.

| ID | Identification days between 2015 to 2019 |
| --- | --- |
| #008 | 46 |
| #014 | 92 |
| #034 | 22 |
| #036 | 58 |
| #045 | 41 |
| #055 | 41 |
| #078 | 35 |
| #182 | 53 |
| #217 | 36 |
| #226 | 51 |
| #231 | 41 |
| #237 | 26 |
| #266 | 49 |
| #306 | 61 |
| #401 | 90 |
| #408 | 42 |
| #452 | 36 |
| #506 | 64 |

Table S3. Details of consortship events.

| Event  number | Date |  | Males | | |  | Females | | |
| --- | --- | --- | --- | --- | --- | --- | --- | --- | --- |
|  |  |  | Dyads | SRI  (5years) | Behavior |  | ID | Reproductive status  at consortship | Birth after  consortship |
| 1 | 2015/06/22 |  | #226 #408 | 0.26 | Capture |  | #025 | Unknown | No |
| 2 | 2015/06/30 |  | #226 #408 | 0.26 | Capture |  | #615 | Not receptive | No |
| 3 | 2015/07/04 |  | #014 #036 | 0.32 | Capture |  | #569 | Receptive | Yes |
| 4 | 2015/10/03 |  | #452 #506 | 0.20 | Capture |  | #558 | Receptive | No |
| 5 | 2016/08/15 |  | #014 #182 | 0.35 | Capture |  | #068 | Receptive | Yes |
| 6 | 2016/08/18 |  | #014 #182 | 0.35 | Capture |  | #068 | - | - |
| 7 | 2017/07/10 |  | #014 #182 | 0.35 | Capture |  | #608 | Receptive | No |
| 8 | 2017/07/25 |  | #237 #306 | 0.15 | Capture |  | #504 | Receptive | No |
| 9 | 2017/08/18 |  | #014 #036 | 0.32 | Capture |  | #314 | Receptive | Yes |
| 10 | 2017/08/28 |  | #237 #401 | 0.15 | Capture |  | #314 | - | - |
| 11 | 2017/09/13 |  | #036 #182 | 0.27 | Capture |  | #314 | - | - |
| 12 | 2018/07/08 |  | #401 #506 | 0.26 | Capture |  | #601 | Receptive | Yes |
| 13 | 2018/07/13 |  | #034 #226 | 0.03 | Capture, Bolt, Aggression |  | #608 | Receptive | Unknown |
| 13 | 2018/07/13 |  | #034 #266 | 0.13 | Capture, Bolt, Aggression |  | #608 | - | - |
| 13 | 2018/07/13 |  | #034 #408 | 0.09 | Capture, Bolt, Aggression |  | #608 | - | - |
| 13 | 2018/07/13 |  | #226 #266 | 0.13 | Capture, Bolt, Aggression |  | #608 | - | - |
| 13 | 2018/07/13 |  | #266 #408 | 0.13 | Capture, Bolt, Aggression |  | #608 | - | - |
| 13 | 2018/07/13 |  | #226 #408 | 0.26 | Capture, Bolt, Aggression |  | #608 | - | - |
| 14 | 2018/07/18 |  | #014 #182 | 0.35 | Capture |  | #161 | Receptive | No |
| 15 | 2019/06/26 |  | #008 #014 | 0.14 | Capture |  | #052 | Unknown | No |
| 16 | 2019/07/05 |  | #045 #231 | 0.36 | Capture |  | #564 | Receptive | Yes |
| 17 | 2019/07/09 |  | #266 #306 | 0.22 | Capture |  | #597 | Receptive | No |
| 18 | 2019/07/10 |  | #036 #182 | 0.27 | Capture |  | #507 | Receptive | Yes |
| 19 | 2019/09/11 |  | #008 #036 | 0.13 | Capture |  | #507 | - | - |
| 19 | 2019/09/11 |  | #008 #182 | 0.13 | Capture |  | #507 | - | - |
| 19 | 2019/09/11 |  | #036 #182 | 0.27 | Capture |  | #507 | - | - |

Table S4. Statistics of the simple ratio indices.

| Analyzed days |  | Simple ratio index | | | | | |
| --- | --- | --- | --- | --- | --- | --- | --- |
|  |  | Social differentiation (SE) | Correlation (SE) | Mean | Observed SD | Expected SD  (mean) | p-value  (one-sided) |
| 221 |  | 0.76 (0.06) | 0.86 (0.03) | 0.08 | 0.07 | 0.06 | <0.01 |

Table S5. Statistics of the half weight indices.

| Survey date |  | Half weight index (HWI) | | | | | |
| --- | --- | --- | --- | --- | --- | --- | --- |
|  |  | Social differentiation (SE) | Correlation (SE) | Mean | Observed SD | Expected SD  (mean) | p-value  (one-sided) |
| 221 |  | 0.70 (0.05) | 0.86 (0.03) | 0.14 | 0.11 | 0.09 | <0.01 |

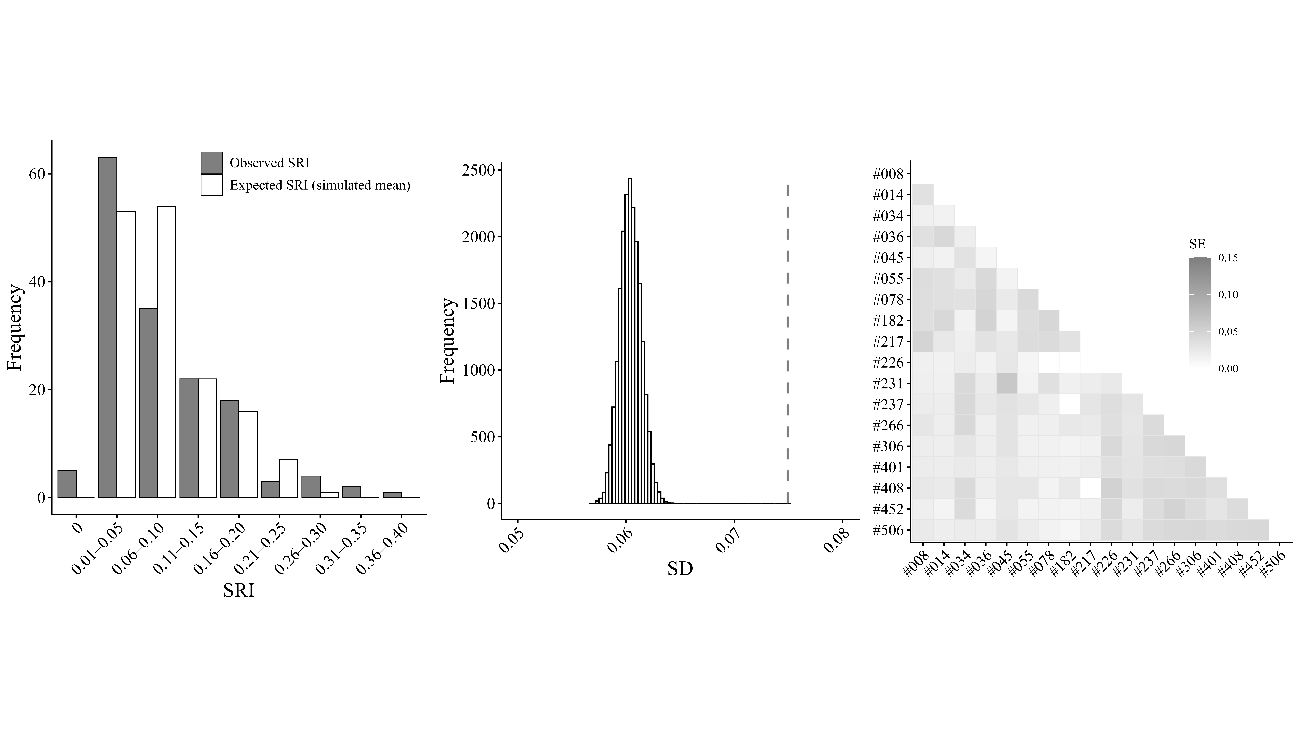
Figure S1. Statistics of the simple ratio indices (SRI). The left panel shows the observed and expected SRI (expected values were calculated by the mean of 20,000 simulations). Center panel shows the observed and expected SD of SRI. The vertical dashed line is the observed value. The right panel shows that SE of SRI among 18 males.

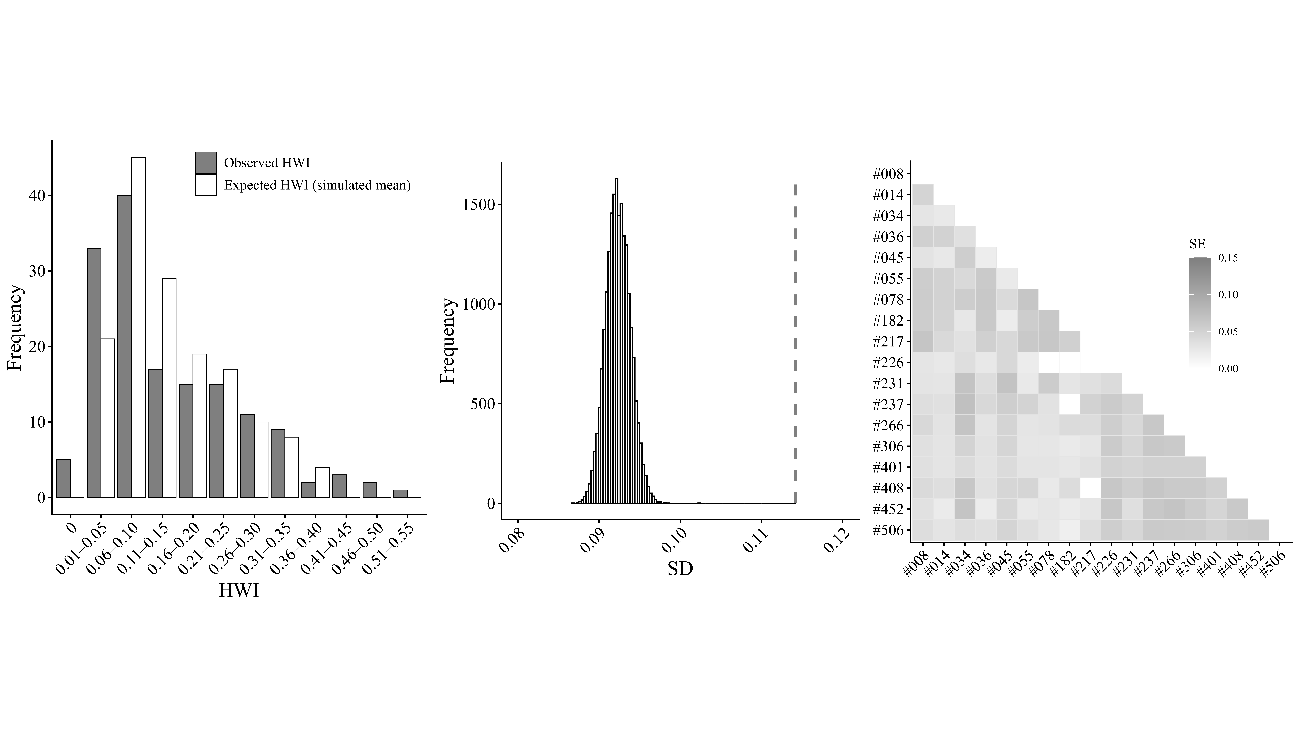

Figure S2. Statistics of the half weight indices (HWI). The left panel shows the observed and expected HWI (expected values were calculated by the mean of 20,000 simulations). Center panel shows the observed and expected SD of HWI. The vertical dashed line is the observed value. The right panel shows that SE of HWI among 18 males.

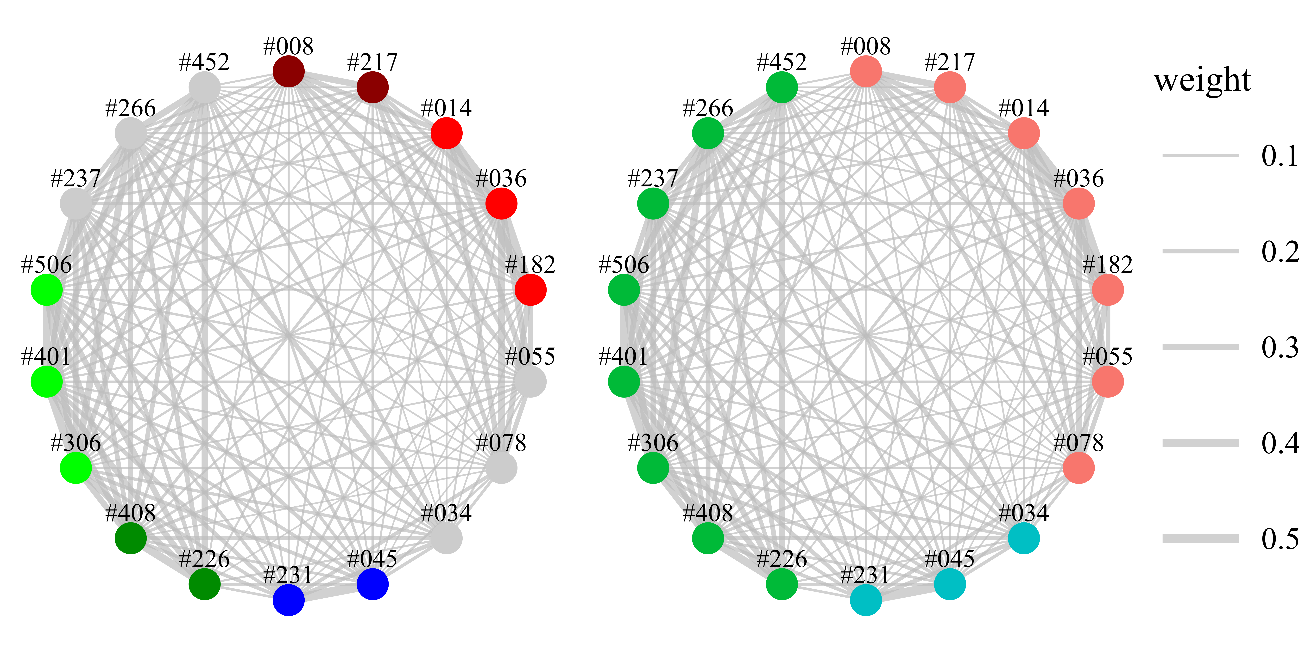

Figure S3. Sociogram constructed by HWI with simple and complex model results (left = simple model, right = complex model). Nodes represent individuals and their color represents unit membership. Note that gray nodes represent individuals that were not assigned to any units. Edges represent association and their width represents association indices.

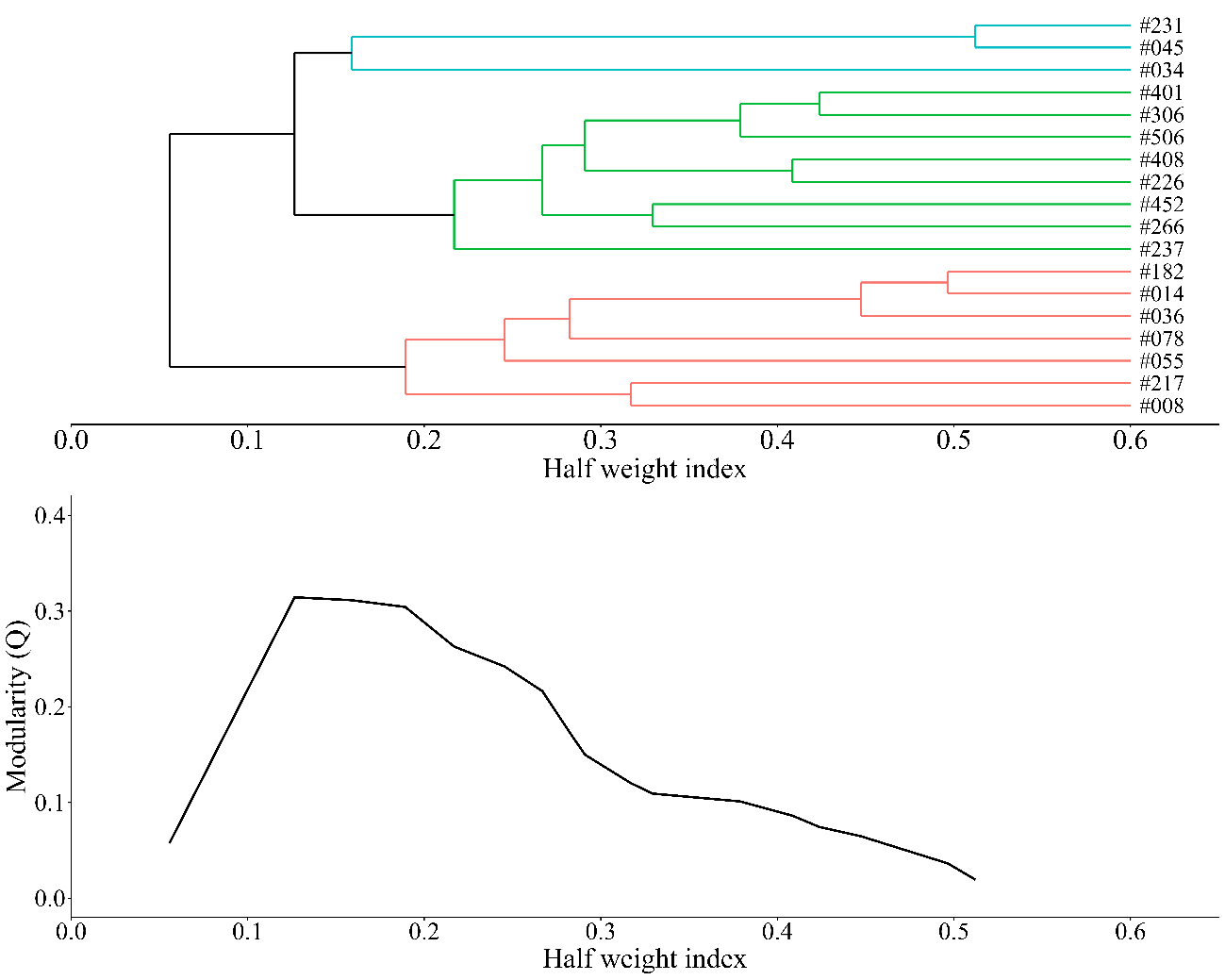

Figure S4. The top dendrogram was constructed by the average-linkage method with HWI. Colors represent units determined by the modularity method. The graph below shows changes in modularity with dendrograms bifurcations.

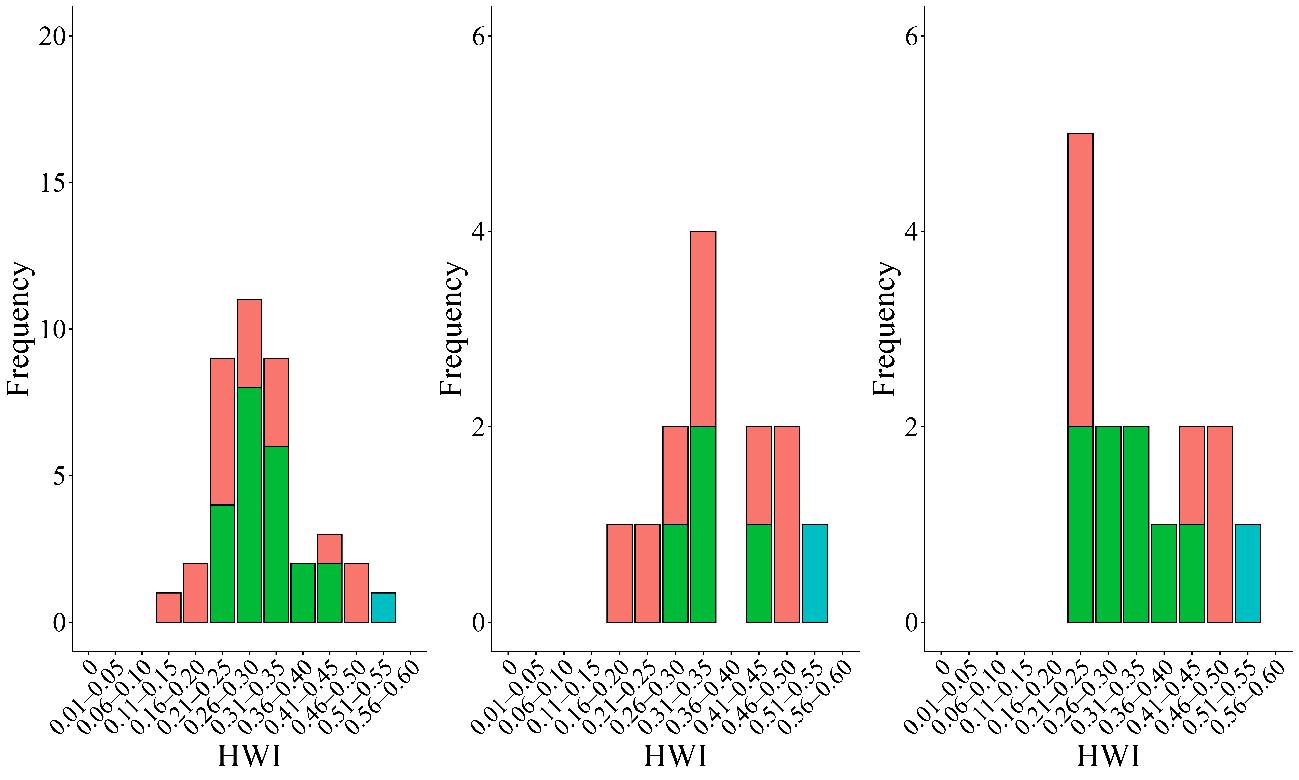

Figure S5. Within unit relation between SRI and behavior. The bar graphs, from left to right, show the results for proximity, rubbing, and consortship. Bar color represents units determined by the complex model (Figure S3 and S4).

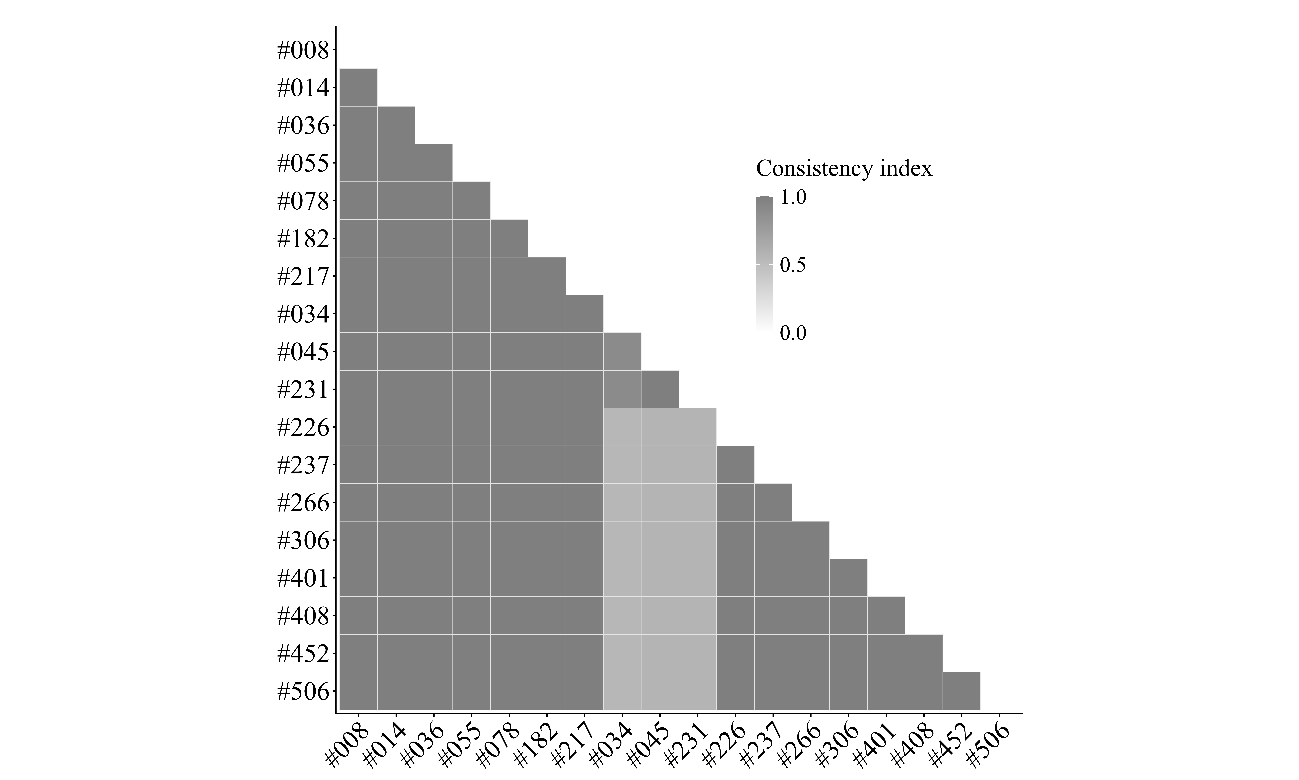

Figure S6. Consistency indices calculated by jackknife simulation with HWI.

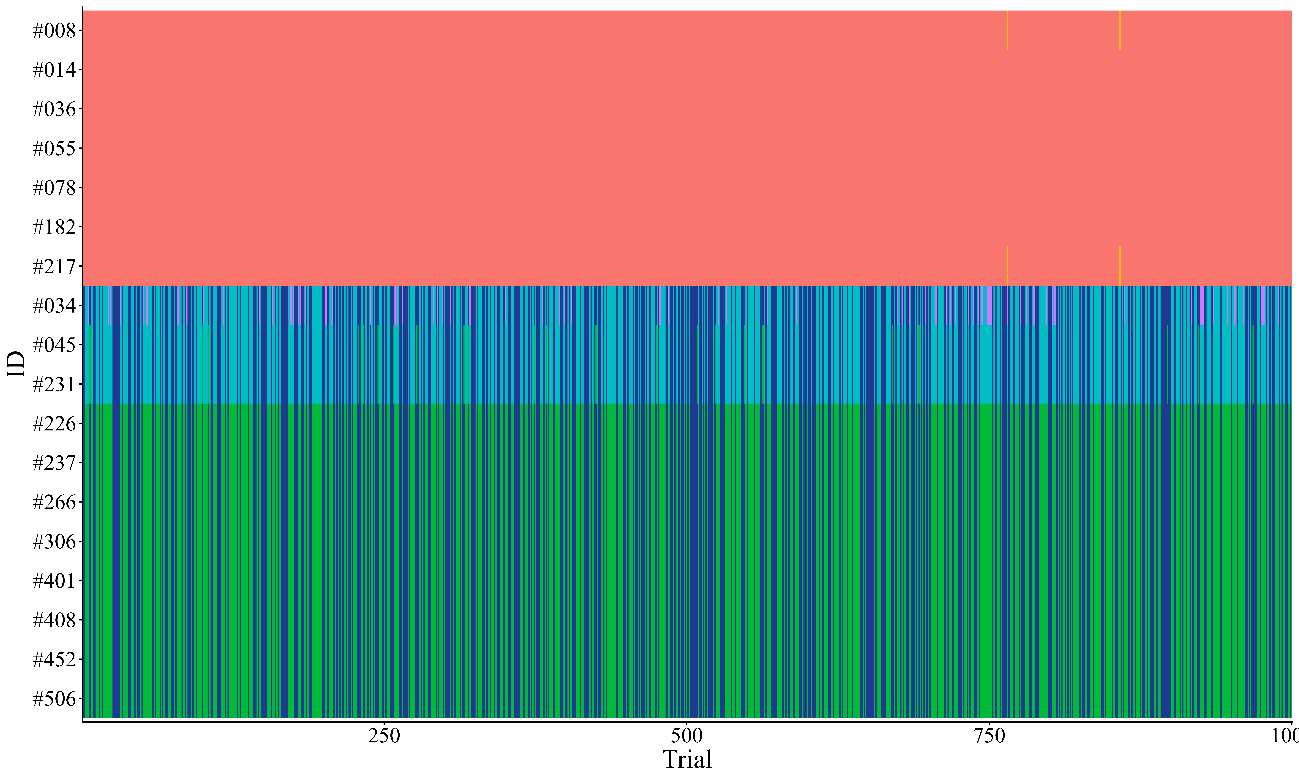

Fig S7. Details of the simulation using HWI. The x-axis represents 1,000 trials. Each color represents unit membership.
